# Pfam domain adaptation profiles reflect plant species’ evolutionary history

**DOI:** 10.1101/2021.07.13.452250

**Authors:** Sarah E. Jensen, Edward S. Buckler

**Affiliations:** School of Integrative Plant Sciences, Plant Breeding and Genetics Division, Cornell University, Ithaca, NY, 14853, USA; Institute for Genomic Diversity, Cornell University, Ithaca, NY, 14853, USA; United States Department of Agriculture, Agricultural Research Service, Ithaca, NY, 14850, USA

## Abstract

The increase in global temperatures predicted by climate change models presents a serious problem for agriculture because high temperatures reduce crop yields. Protein biochemistry is at the core of plant heat stress response, and understanding the interactions between protein biochemistry and temperature will be key to developing heat-tolerant crop varieties. Current experimental studies of proteome-wide plant thermostability are limited by the complexity of plant proteomes: evaluating function for thousands of proteins across a variety of temperatures is simply not feasible with existing technologies. In this paper, we use homologous prokaryote sequences to predict plant Pfam temperature adaptation and gain insights into how thermostability varies across the proteome for three species: maize, Arabidopsis, and poplar. We find that patterns of Pfam domain adaptation across organelles are consistent and highly significant between species, with cytosolic proteins having the largest range of predicted Pfam stabilities and a long tail of highly-stable ribosomal proteins. Pfam adaptation in leaf and root organs varies between species, and maize root proteins have more low-temperature Pfam domains than do Arabidopsis or poplar root proteins. Both poplar and maize populations have an excess of low-temperature mutations in Pfam domains, but only the mutations identified in poplar accessions have a negative effect on Pfam temperature adaptation overall. These Pfam domain adaptation profiles provide insight into how different plant structures adapt to their surrounding environment and can help inform breeding or protein editing strategies to produce heat-tolerant crops.

## INTRODUCTION

Understanding the interactions between temperature and plant development is becoming more important as global temperatures rise. Temperature-induced abiotic stress can affect every aspect of plant development, from morphology and physiology to biochemistry. Climate change is predicted to increase the levels of extreme temperature stress to which plants are exposed, and as a result, shift existing ranges and alter planting times for important agronomic species[1–3]. An increase in growing season temperatures will affect both crops and natural vegetation. In crops, increasing temperatures are predicted to substantially reduce yields, with estimates of 1-7% yield reduction in major crop species for every 1℃ global temperature increase [4]. In other species, increases in maximum annual temperature are expected to increase the rate of local extinction and threaten both plant biodiversity and larger ecological networks that rely on keystone plant species [5,6]. Preventing substantial decreases in crop yield and ecological disasters due to species extinction requires a better understanding of how plant species adapt to high-temperature environments.

Specific metabolic pathways involved in plant temperature sensing and response, including thermomorphogenesis, have been well-characterized [7]. Less-well studied is how the temperature of the environment shapes molecular evolution across the proteome. Interactions between biochemistry and temperature affect DNA, RNA, and protein composition, and are consistent across species and even across phylogenetic domains [8–12]. In plants, genome size, GC content, and proteome composition have been correlated with environmental temperature [13,14]. Temperature is also correlated with changes in diverse cellular processes including chromatin remodeling, lipid membrane composition, photosynthetic capacity, and hormone signaling in addition to protein expression [12,15–18].

Because temperature affects protein folding, activity, and function, it likely also influences protein shifts toward increased or decreased stability [19–22]. Some proteins, like Phytochrome B and ELF3, are thermosensor proteins that have evolved to be only marginally stable. Denaturing these proteins initiates important stress response and circadian signaling pathways [23,24]. Other proteins remain only marginally stable because negative selection cannot prevent accumulation of destabilizing mutations. Purifying selection acts to avoid complete protein unfolding, but is too weak to optimize stability for most proteins [25]. As a result, there is a distribution of protein stabilities across the proteome. Most proteins are stable, but a significant proportion of the proteome - perhaps as much as 10-15% of all proteins - are only minimally stable and can be denatured by temperature shifts of as little as 4℃ [26]. Plants experience temperature shifts of this magnitude or larger daily. It remains unclear how well plant proteomes will be able to adapt to the increasing global temperatures projected in the next century.

Proteins are composed of functional regions, or domains, that carry out specific biochemical reactions. We hypothesize that Pfam domain temperature adaptation affects plant thermotolerance and that Pfam adaptation profiles differ across species. Because biochemical and physical constraints on protein function are similar across all species, we use comparisons to prokaryotic Pfam domains to create profiles of temperature adaptation for maize, Arabidopsis, and poplar [27]. We compare Pfam adaptation profiles between plant organs and organelles and look at mutation effects across populations to investigate how amino acid mutations affect adaptation profiles within a species. These estimates of protein stability provide insight into how plant proteomes evolve and will be a useful starting point from which to develop strategies to improve plant heat tolerance as global temperatures rise.

## MATERIALS AND METHODS

### Identifying eukaryote Pfam domains

Pfam domains were identified in maize (AGPv3; [28]), Arabidopsis (TAIR10; [29]), and poplar (v3.0; [30]) reference proteomes using the hmmscan function in HMMER3 and default parameters [31]. Pfam domains were aligned to existing prokaryote Pfam domain alignments, numerically recoded to reflect amino acid physicochemical properties, and assigned Pfam domain clusters based on sequence similarity to the prokaryotic sequences in the alignment. Realignment utilized the mafft --add function with default parameters to maintain original alignment coordinates [32]. Prokaryote GWAS results from Jensen et al. (*in prep*) were used to identify Pfam domain positions significantly associated with temperature. Each aligned amino acid residue in the maize, Arabidopsis, and poplar Pfam domains were compared to the prokaryote Pfam domain GWAS results to estimate temperature adaptation at that residue. Positions that were not significantly associated with temperature in prokaryotes were removed.

### Population nonsynonymous mutations

Previously-published VCF files containing sequence data and collection locations were downloaded for 1119 Arabidopsis [33], 31 maize [34], and 567 poplar accessions [35,36]. Nonsynonymous mutations were identified with snpEff, and the ‘-proteinFasta’ option was used to output both reference and variant protein fasta files for each accession [37]. SnpEff output files were filtered to identify unique nonsynonymous mutations in each accession. Allele frequencies were calculated from the original VCF files for all biallelic sites using vcftools [38]. Each identified nonsynonymous mutation and the corresponding reference amino acid was recorded, as was the mutation position and allele frequency.

### Calculating mutation temperature sensitivity

Average residue temperature was calculated by averaging the optimal growth temperatures (OGTs) across all prokaryotes sharing an amino acid at that site. Residue temperature averages were calculated for all positions that were significantly associated with OGT in prokaryotes (prokaryote OGT values range from 10.1-96.3ºC) and were recorded for all identified Pfam domains in maize, Arabidopsis, or poplar. These residue temperatures were used as a thermal proteome profile for each species based on the reference proteome (Figure 1). For sites with nonsynonymous mutations identified by snpEff, optimal temperature values were recorded for both the reference amino acid and the variant amino acid. Mutation effects were compared between the major allele amino acid and the minor allele amino acid. For each site with a nonsynonymous mutation, the average prokaryote OGT for the ancestral amino acid was compared to the average prokaryote OGT for the derived amino acid to see whether the mutation likely increased or decreased Pfam domain thermostability. By necessity, this analysis focuses on single residues in isolation and does not consider interactions between amino acids within a folded protein.

**Figure 1:**
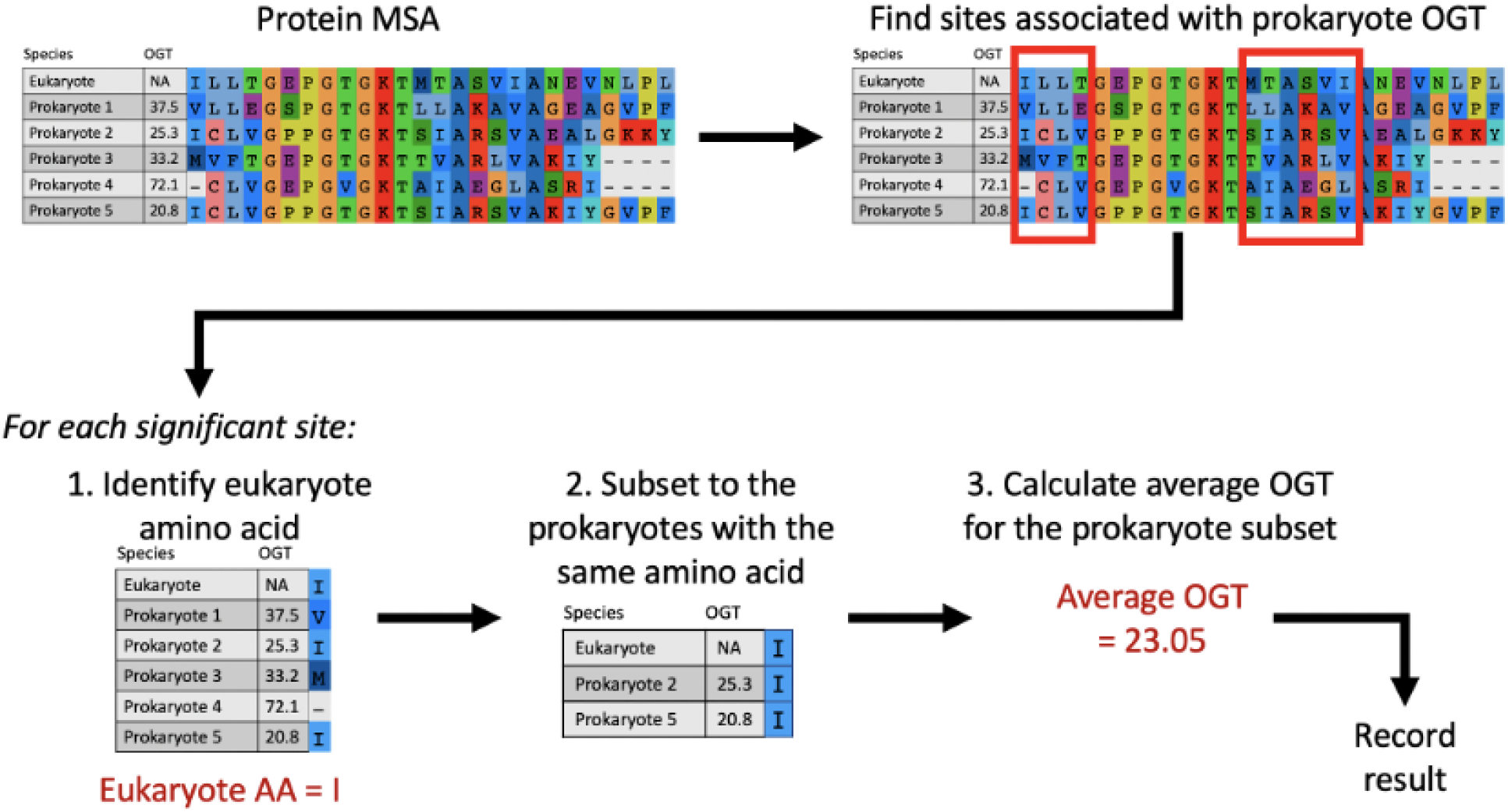
Process for calculating temperature sensitivity of each amino acid residue. Briefly, Pfam domain observations from maize, Arabidopsis, or poplar are aligned to an existing prokaryote multiple sequence alignment. Only the subset of sites that are significantly associated with prokaryotic optimal growth temperature (OGT) are kept. At each significant site, the eukaryote amino acid is identified, and the average OGT for prokaryotes sharing the same amino acid is recorded as the ‘optimal’ temperature of that residue and position.

### Tissue- and organelle-specific protein expression

Organ-specific protein expression for Col-0, B73, and Nisqually-1 were identified from protein and mRNA expression atlas datasets [39–42]. Organelle expression data for plastid, mitochondria, and cytosol proteins was obtained from the Plant Proteome Database [43] for maize and the SUBA database [44] for Arabidopsis. Arabidopsis proteins were assigned a single unique organellar location using SUBAcon organelle calls [45].

Reference proteins were classified as leaf-expressed proteins or root-expressed proteins according to protein or mRNA expression levels in each organ. If multiple samples from leaf or root tissues were available, then these were merged to create a larger subset of proteins for testing. For maize and Arabidopsis, only proteins that were uniquely expressed in one organ and not the other were kept for the analysis. In poplar, the mRNA expression dataset contained too few organ-specific genes for comparison. Instead, the top 10% of mRNA transcripts were identified in each organ and used to compare protein stability. Protein counts for each organ and organelle are listed in Table 1. Pfam adaptation distributions were compared between organelles and between leaves and roots.

**Table 1:** Gene counts for maize, Arabidopsis, and poplar in each organ and organelle.

|  | Number of unique proteins |  |  |
| --- | --- | --- | --- |
|  | Maize | Arabidopsis | Poplar |
| Leaf-specific | 933 | 346 | 730 |
| Root-specific | 529 | 864 | 560 |
| Cytosol | 254 | 2053 | - |
| Ribosome | 208 | 239 | - |
| Plastid | 1375 | 1357 | - |
| Mitochondria | 155 | 735 | - |

### GO analysis

Gene ontologies associated with genes in Arabidopsis thaliana or Zea mays were used to identify terms enriched in proteins with high temperature stability estimates in organellar proteins [46,47]. The topGO R package was used to compare GO terms for protein with average predicted-stability values above 40℃ to the set of all proteins with predicted stabilities [48]. topGO was also used to evaluate the trimodal distribution of root Pfam domain adaptation, with low-temperature domains considered to have values < 30.4℃, moderate-temperature domains having values from 30.4-34.4℃, and high-temperature domains having values > 34.4℃.

## RESULTS

### Predicted Pfam adaptation correlates with half-life and expression measurements

Protein expression and half-life experiments in Arabidopsis were used to compare Predicted Pfam Adaptation (PPA) from prokaryote-based estimates of Pfam domain adaptation to experimental values for rosette leaf protein expression (3618 proteins) and protein half-life (750 proteins) [39,49]. There is a weak positive correlation between log-linearized protein half-life and PPA (Figure 2A), and also between PPA and normalized protein expression values (Figure 2B). Surprisingly, the correlation between measured protein half-life and measured protein expression is even weaker than the correlations with Pfam adaptation estimates (Figure 2C).

**Figure 2:**
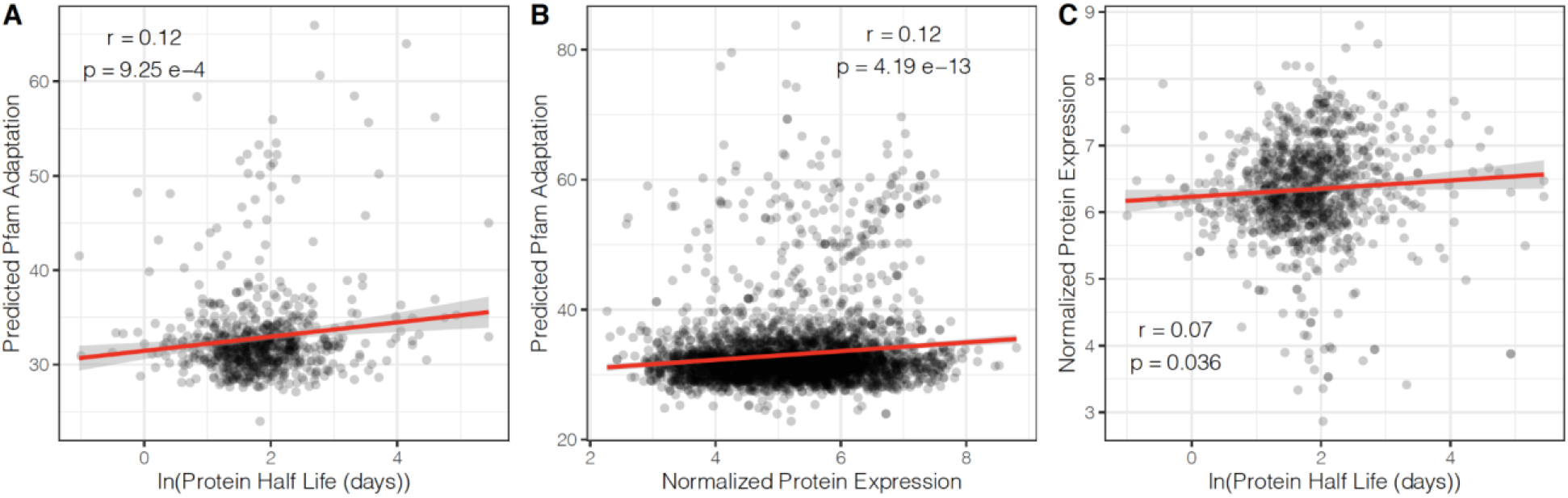
OGT-based predicted Pfam adaptation (PPA) values are weakly correlated with measured expression and stability values. A) correlation between predicted protein stability and protein half-life (r = 0.12, p = 9.25e-4, n = 750); B) correlation between predicted stability and protein expression (r = 0.12, p = 4.19e-13, n = 3618); C) correlation between protein half-life and expression level (r = 0.07, p = 0.036, n = 1001).

### Predicted Pfam adaptation differs between organs and organelles

Plant organs function in soil and air temperatures that fluctuate daily. In addition to changing air temperatures, leaf proteins experience variable light intensities and qualities, while root proteins function in the context of the larger rhizosphere. We hypothesized that the differences in environments experienced by these different plant organs would lead to different PPA profiles between Pfam domains in root-expressed proteins and leaf-expressed proteins. Organ-specific protein expression data are available for maize and Arabidopsis reference genomes, and organ-specific mRNA datasets are available in poplar [39,41,42]. These data were used to determine whether protein stability differs between plant leaves and roots. In maize there is a significant difference in average root protein stability and average leaf protein stability, with a significantly higher average stability for leaf proteins (t-test, p = 0.0056; Figure 3A, Supplemental Figure 1). Neither Arabidopsis nor poplar have tissue-specific differences in protein stability between leaf and root proteins as predicted by the OGT pipeline (t-test, p > 0.05; Figure 3B-C). In all three species leaf proteins have higher average expression than root proteins, so the stability difference observed in maize proteins is not likely to be driven solely by expression differences between the tissues (Supplemental Figure 2).

**Figure 3:**
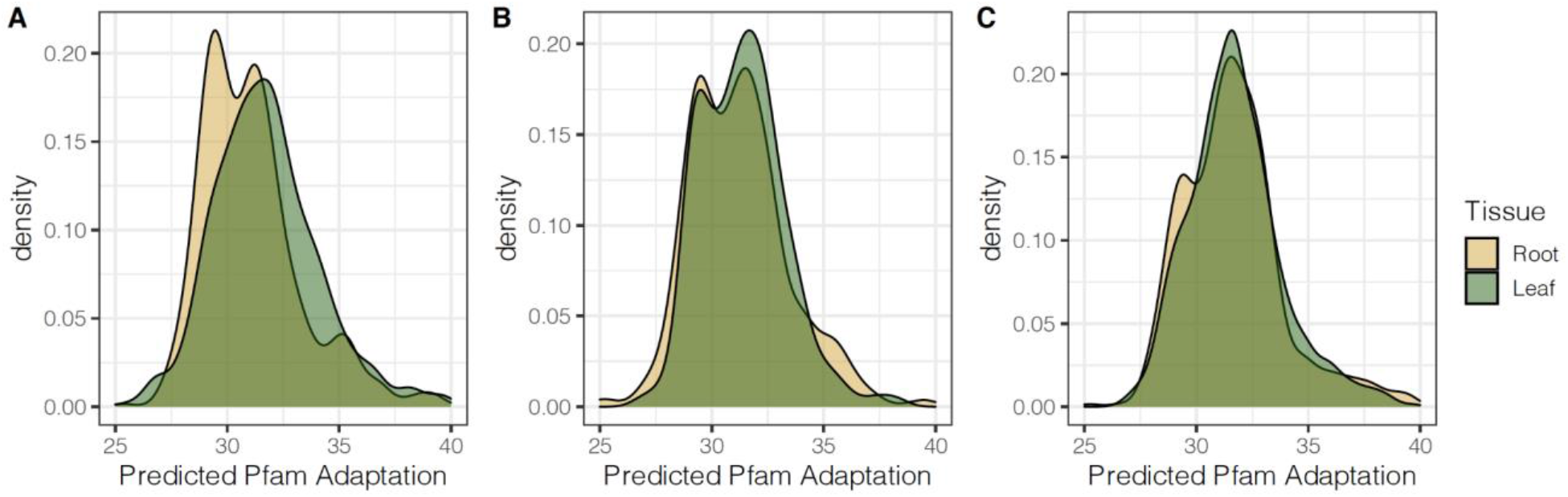
Leaf and root protein adaptation distributions predicted by the prokaryote optimal temperature pipeline. Organs have significantly different temperature profiles in maize (A) but not in Arabidopsis (B) or poplar (C). For clarity, distributions are truncated at 40℃, but all three have tails that extend beyond 60℃. Full distributions are included as Supplemental Figure 1.

Maize root Pfam domains separated into a trimodal distribution, with a low PPA (< 30.4℃), moderate PPA (30.4-34.4℃), and high PPA (>34.4℃) groups of Pfam domains. GO enrichment analysis showed significant enrichment for specific classes of proteins in these three groups. Low-PPA domains were enriched for GO terms related to ion transport, vacuolar structures, and enzymes involved in redox reactions, including antioxidant, peroxidase, oxidoreductase, and hydrolase GO terms. Moderate-PPA domains were enriched for basic cell processes including transport and signal transduction, membrane structures, and kinase, transferase, and peptidase enzymes. High-PPA domains seemed to contain a mix of proteins, with enrichment for GO terms related to binding processes and negative regulation (Supplemental Table 1).

In endotherms, respiratory chain and mitochondrial proteins operate at higher temperatures than normal body temperature, which suggests that some organelles operate at higher-than-ambient temperatures relative to the rest of the cell [50,51]. PPA comparisons across plant organelles suggests that organelles also operate at different temperatures within plant cells. Protein subcellular localization data are available for both Arabidopsis and maize [43,44]. PPA differs between organelles, and patterns of organellar stability are consistent in both maize and Arabidopsis. A Games-Howell test, which is robust when variances are unequal between groups, was used to compare protein stability predictions across organelles. Cytosol, plastid, and mitochondrial proteins have significantly different average stabilities (p < 0.0001 for all pairwise comparisons, Games-Howell test; Table 2). PPA varies widely within each organelle, and further investigation shows that ribosome and translation-associated GO terms are enriched in the set of Arabidopsis proteins with highest PPA values suggesting that proteins involved in translation have higher-than-average Pfam stability (Fisher’s test, p < 0.0001; Supplemental Table 1). GO enrichment analysis for maize did not indicate enrichment in ribosomal proteins, but GO terms for non-membrane-bounded organelles and for protein-containing complexes were enriched (Figure 4A, Supplemental Table 2). Ribosomal proteins are plotted separately from other organelles for both maize and Arabidopsis. A long tail of stable cytosolic proteins remains in Arabidopsis, suggesting that many other cytosolic proteins are also adapted to high temperatures. There is also a significant difference between plastid and mitochondrial PPA values, with plastid proteins predicted to be adapted to higher temperatures than mitochondrial proteins (Figure 4).

**Table 2:** Games-Howell pairwise comparisons of predicted protein stabilities across organelles in maize and Arabidopsis.

| Species | Organelle 1 | Organelle 2 | p-value (adj.) |
| --- | --- | --- | --- |
| Maize | Cytosol | Ribosome | 8.71E-13 |
| Maize | Cytosol | Plastid | 9.20E-14 |
| Maize | Cytosol | Mitochondria | 2.65E-10 |
| Maize | Ribosome | Plastid | 4.45E-6 |
| Maize | Ribosome | Mitochondria | 4.19E-14 |
| Maize | Plastid | Mitochondria | 0 |
| Arabidopsis | Cytosol | Ribosome | 7.74E-14 |
| Arabidopsis | Cytosol | Plastid | 0 |
| Arabidopsis | Cytosol | Mitochondria | 0 |
| Arabidopsis | Ribosome | Plastid | 0 |
| Arabidopsis | Ribosome | Mitochondria | 0 |
| Arabidopsis | Plastid | Mitochondria | 4.57E-05 |

**Figure 4:**
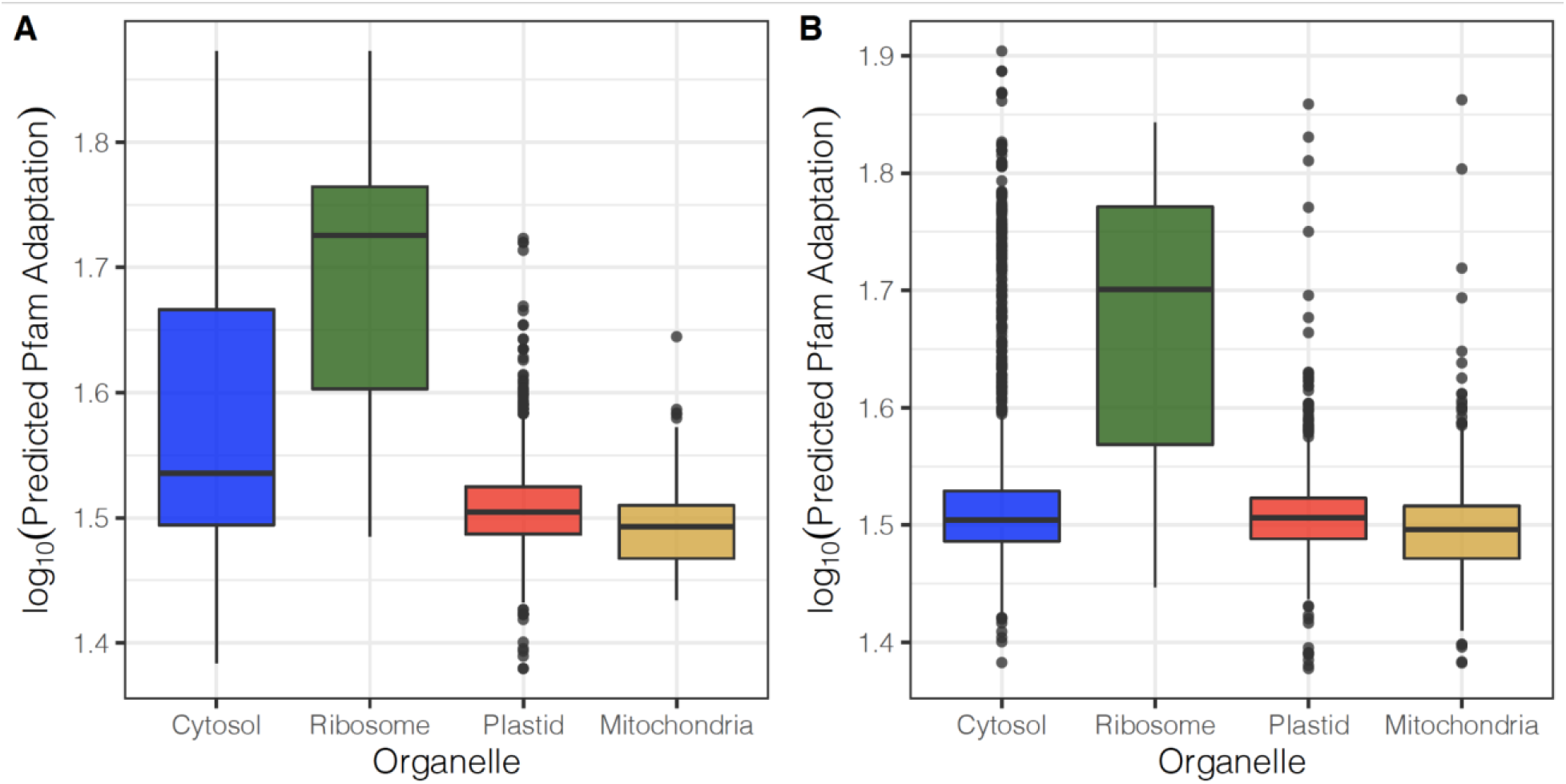
Predicted Pfam adaptation distributions differ in cytosol, ribosome, mitochondria, and plastid organelles in maize (A) and Arabidopsis (B). Group means were compared with a one-way ANOVA with unequal variances and pairwise comparisons were made using a Games-Howell test.

### Cumulative mutation effects vary within a population

Many plant species extend across a wide range of environments. Maize accessions, for example, grow in a range of environments, from cool, high-altitude environments to tropical environments [15]. Amino acid mutations that accumulate as a species expands its range may reflect adaptation to new environments, so we hypothesized that nonsynonymous mutations would affect Pfam adaptation predictions and be related to the environment from which an accession is collected. To test this hypothesis, we identified nonsynonymous mutations in populations of maize, Arabidopsis, and poplar and determined whether the mutation increased or decreased PPA relative to the major allele. Maize shows a particularly interesting pattern with this analysis: individual amino acid mutations in maize tend to decrease PPA and reduce Pfam adaptation to high temperature. However, the cumulative effects of maize mutations tend to increase PPA overall (Figure 5A, 5D). Arabidopsis and poplar show more expected distributions, with a consistent pattern between the proportion of mutations that decrease PPA and the overall effect of those variants. Arabidopsis accessions tend to accumulate mutations that increase PPA across the proteome (Figure 5B, 5E), while poplar accessions tend to accumulate mutations that decrease PPA (Figure 5C, 5F).

**Figure 5:**
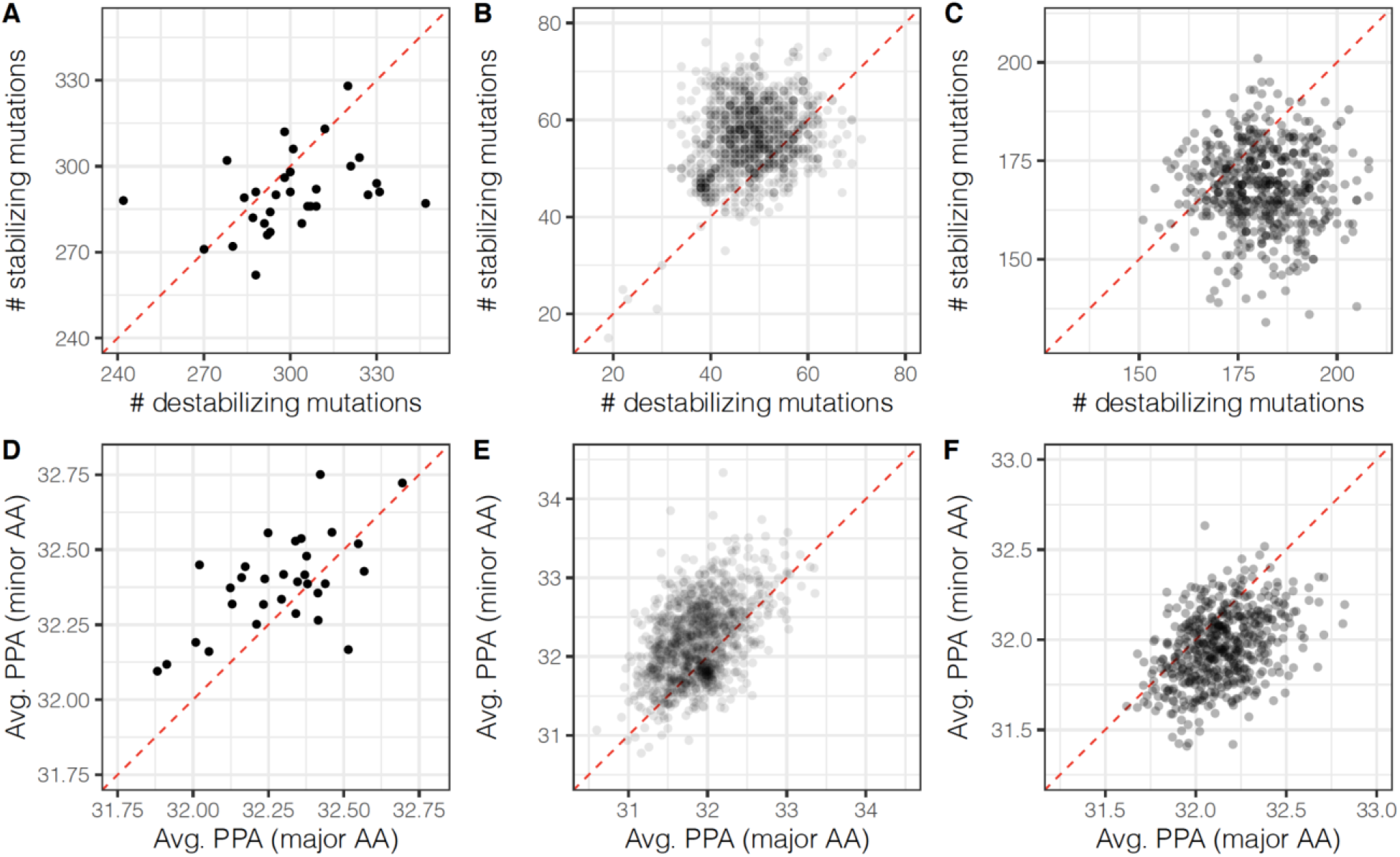
Predicted amino acid mutation counts and stability effects are consistent in Arabidopsis and poplar, but not in maize. Each point in the plots shows the cumulative count (A-C) or effect (D-F) of nonsynonymous amino acids for a single landrace. Maize tends to accumulate individual mutations that reduce PPA (A), but these mutations have an overall positive effect on PPA (D). Arabidopsis accessions accumulate individual mutations that increase PPA (B) and these mutations also have a positive cumulative effect on PPA (E). Poplar accessions accumulate individual mutations that reduce PPA (C) and the overall effect of these mutations is reduced PPA (F). In A-C, the red dotted line indicates the point where the number of mutations that increase PPA equals the number of mutations that decrease PPA. In D-F the red dotted line indicates the point where the cumulative effects of nonsynonymous mutations is zero.

Maize originates from the Balsas River Valley in Mexico [52] and Arabidopsis is thought to originate from Morocco [53]. Poplar samples in the dataset come from the American Pacific Northwest coast [42]. To see whether mutations have a different effect as a species expands beyond its center of origin we calculated the difference between PPA estimates for the major allele and the minor allele and summed the effects across all nonsynonymous mutations in the individual. In all three species there is a negative relationship between the net mutation effects and the distance from the center of origin suggesting that the cumulative effect of mutations become more destabilizing as a species expands (Figure 6).

**Figure 6:**
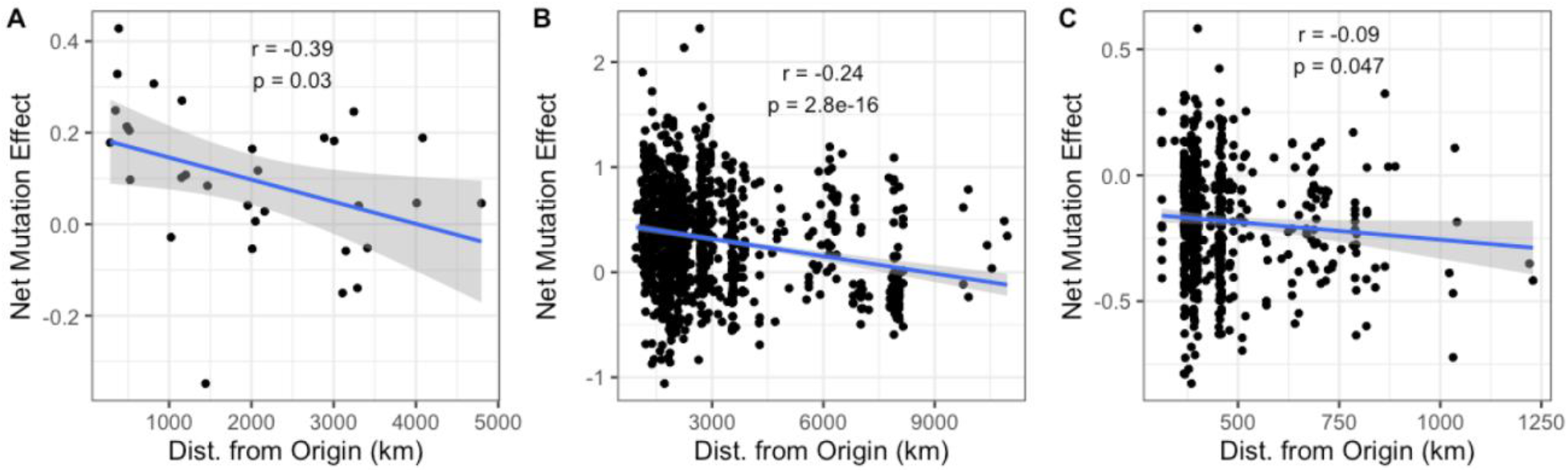
Net effects of mutations on predicted Pfam adaptation (major allele PPA – minor allele PPA) reduce temperature adaptation as accessions move further from the center of origin for A) maize, B) Arabidopsis, and C) poplar. Distance from the center of origin was determined with the geopy python package. The estimated center of origin for the species was (17.9373, − 102.1360) for maize, (31.7917, 7.0926) for Arabidopsis, and (45.5001, −118.0013) for poplar based on the center of origins identified for each species.

## DISCUSSION

We predicted Pfam adaptation across Arabidopsis, maize, and poplar populations to see if protein adaptation is distributed differently across three species with different growth strategies. Because proteins are energetically expensive cell components, the translational robustness hypothesis predicts that highly abundant proteins in the cell will be more stable, as will proteins with longer half-life [49,54,55]. Consistent with this hypothesis, we find that our predicted Pfam adaptation values are positively and modestly correlated with both protein half-life and protein expression. One benefit of using protein PPA estimates over measured stability values is that these estimates can be calculated for any species and can capture a larger proportion of the proteome than experimental studies.

Proteins should be exquisitely adapted to their environment, but local environments may differ between different parts of a plant. In maize, leaf-expressed proteins are adapted to higher temperatures than root-expressed proteins, but there is no difference in PPA between leaf and root proteins in Arabidopsis or poplar. This difference in organ effect across species is unlikely to be due to differences in protein expression between leaf and root tissues because all three species had higher protein expression in leaves, but only maize shows a difference in thermal adaptation profile. Interestingly, the difference in maize appears to be a result of higher PPA values in maize leaf proteins relative to root proteins. Unlike both Arabidopsis and poplar, maize is a C4 grass species adapted to grow in hot environments, and maize net photosynthesis is maximized between 30-35℃ [56]. In contrast, net photosynthesis reaches a maximum in poplar between 25-30℃ [57] and Arabidopsis maximum photosynthetic rate occurs around 25℃ [58]. We hypothesize that the observed difference in leaf and root protein temperature profiles in maize reflects the higher temperatures in which this species is photosynthetically active.

Consistent with observations in birds and mammals, maize and Arabidopsis show similar protein stability profiles across organelles, and the observed PPA distributions are also consistent with previous work comparing protein half-life and turnover rates in Arabidopsis [49–51]. Surprisingly, cytosolic proteins showed a wide distribution of predicted Pfam adaptation values, with a long tail of high-PPA proteins that are enriched for ribosome and cytosolic ribosome GO terms. Ribosomal proteins are expressed at high levels in nearly every cell, and their high expression levels in addition to their importance for translation may explain their shifted stability distributions relative to other proteins in the cytosol [59].

To see how amino acid mutations affect predicted Pfam adaptation, we compared mutations across multiple accessions of maize, Arabidopsis, and poplar. Most nonsynonymous mutations in maize and poplar decrease PPA relative to the major allele, while Arabidopsis amino acid mutations increase PPA relative to the major allele. Intriguingly, the maize accessions used in this study have a tendency to accumulate destabilizing PPA mutations, but the net effect of those mutations tends to be increased adaptation to high temperatures. This pattern suggests that there are many weakly destabilizing mutations in maize that lower temperature adaptation, but whose effects can be offset by a few mutations that substantially increase thermal stability. The extent to which these mutations affect protein function should be a topic for future studies.

The last glacial maximum occurred only 18,000 years ago; a relatively short time in the context of plant evolutionary history. At that time, global temperatures were 4-8℃ lower than modern temperatures, even in the tropics [60]. Arabidopsis and maize are both annual pioneer species that expanded throughout the world within the last 4,000-10,000 years as vegetation patterns changed in response to temperature increases following the end of the glacial period [61–63]. Both species also benefited from large effective population sizes during their expansion [53,64]. We hypothesize that short generation times and large effective population sizes allow pioneer species to evolve and expand rapidly, leaving signatures of directional evolution within the proteome. Unlike maize and Arabidopsis, poplar is a long-lived perennial species. Like other trees, it may experience an adaptational lag that limits its ability to adapt to current climate conditions [65]. We expect allelic diversity to decline and deleterious mutations to accumulate as species expand to new environments far from the center of origin [34,53]. Our observed negative correlation between net mutation effect and distance from origin in all three species is consistent with this expectation, and the weaker relationship in poplar may stem from the slow rate of molecular evolution observed in trees [66].

Proteins that are only minimally stable are of particular interest for understanding plant heat tolerance because temperature sensitivity has been linked to a loss of specific important proteins that disrupt cell function and lead to cell death [54]. The results presented here demonstrate that protein thermostability profiles differ across organelles, and to some extent across tissues, and suggest that population-wide mutation effects also differ across species. Importantly, these results show that even successful pioneer species may be only marginally successful at avoiding destabilizing protein mutations, as shown by the large number of destabilizing mutation effects observed in maize. This suggests that targeted human interventions will be needed to help adapt crops and wild species to higher average temperatures. Further studies are needed to understand the complex interactions between protein thermostability and plant heat stress tolerance. Maintaining crop yields and mitigating ecological disaster due to local species extinctions will likely require both intensive breeding for heat tolerant varieties and targeted genome editing to stabilize plant proteins.

## SUPPORTING INFORMATION

Scripts used for the analyses described can be found on Bitbucket at bitbucket.org/bucklerlab/proteomethermalprofiling/. Data files can be found on CyVerse Data Commons at /iplant/home/shared/commons_repo/curated/Jensen_plantProteomeProfiling_Jun2021.

