## Supplemental Figures 1-2, Supplemental Tables 1-2 for "Pfam domain adaptation profiles reflect plant species’ evolutionary history"

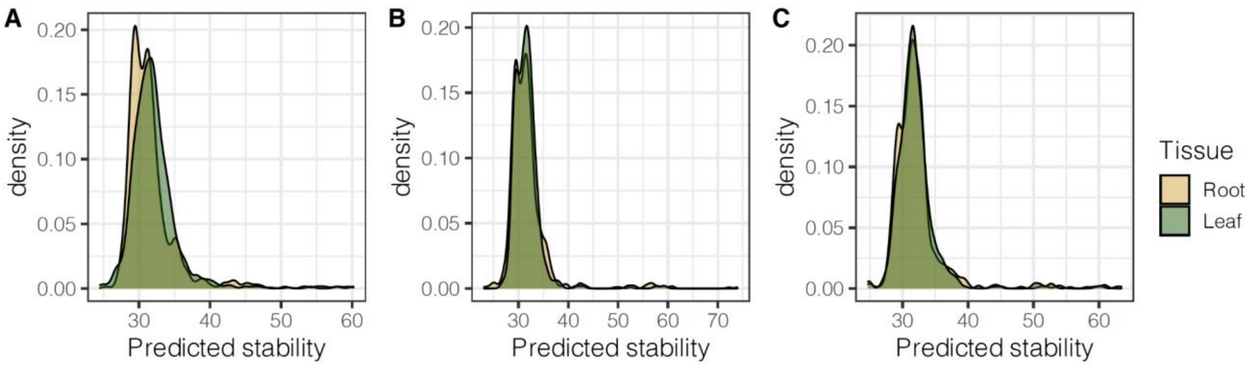


Supplemental Figure 1: Full leaf and root protein stability distributions predicted by the prokaryote optimal temperature pipeline in A) maize, B) Arabidopsis, and C) poplar.

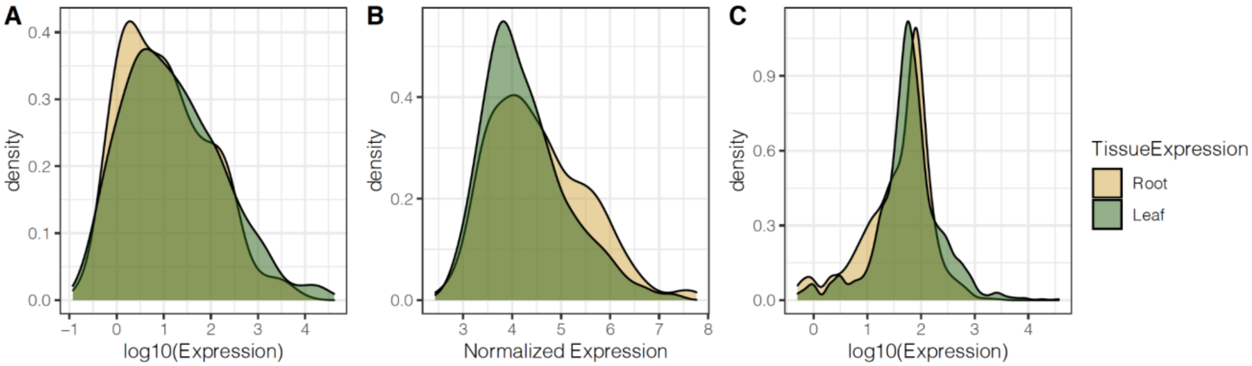


Supplemental Figure 2: Leaf and root protein expression distributions for A) maize, B) Arabidopsis, and C) poplar. In all three species leaf proteins are expressed at higher levels than root proteins. Maize: p=0.02, Arabidopsis: p=9.1E-06, Poplar: p=8.3E-13.

Supplemental Table 1: GO term enrichment for Arabidopsis Pfam domains with high PPA (PPA > 40). Bolded p-values pass a 5% Bonferroni multiple testing correction. Only the top 30 GO terms are shown.

| GO ID | Term | Annotated | Significant | Expected | p-value |
|---|---|---|---|---|---|
| GO:0022626 | cytosolic ribosome | 64 | 49 | 4.12 | **< 1e-30** |
| GO:0005840 | ribosome | 81 | 53 | 5.22 | **< 1e-30** |
| GO:0043228 | non-membrane-bounded organelle | 131 | 61 | 8.44 | **< 1e-30** |

| | | | | | |
|---|---|---|---|---|---|
| GO:0043232 | intracellular non-membrane-bounded organelle | 131 | 61 | 8.44 | **< 1e-30** |
| GO:1990904 | ribonucleoprotein complex | 90 | 51 | 5.8 | **< 1e-30** |
| GO:0032991 | protein-containing complex | 188 | 70 | 12.11 | **< 1e-30** |
| GO:0044391 | ribosomal subunit | 59 | 40 | 3.8 | **< 1e-30** |
| GO:0005829 | cytosol | 186 | 61 | 11.98 | **6.80E-30** |
| GO:0022627 | cytosolic small ribosomal subunit | 25 | 23 | 1.61 | **3.50E-26** |
| GO:0015935 | small ribosomal subunit | 30 | 25 | 1.93 | **4.70E-26** |
| GO:0022625 | cytosolic large ribosomal subunit | 19 | 14 | 1.22 | **1.20E-13** |
| GO:0015934 | large ribosomal subunit | 29 | 15 | 1.87 | **3.00E-11** |
| GO:0032993 | protein-DNA complex | 5 | 5 | 0.32 | 1.10E-06 |
| GO:0005622 | intracellular | 2620 | 192 | 168.76 | 3.10E-05 |
| GO:0005634 | nucleus | 589 | 61 | 37.94 | 4.10E-05 |
| GO:0000428 | DNA-directed RNA polymerase complex | 6 | 4 | 0.39 | 0.00023 |
| GO:0030880 | RNA polymerase complex | 6 | 4 | 0.39 | 0.00023 |
| GO:0055029 | nuclear DNA-directed RNA polymerase complex | 6 | 4 | 0.39 | 0.00023 |
| GO:1902494 | catalytic complex | 58 | 12 | 3.74 | 0.00024 |
| GO:0005681 | spliceosomal complex | 11 | 5 | 0.71 | 0.00036 |
| GO:1990234 | transferase complex | 24 | 7 | 1.55 | 0.00056 |
| GO:0061695 | transferase complex, transferring phosphatase | 8 | 4 | 0.52 | 0.00095 |
| GO:1905368 | peptidase complex | 5 | 3 | 0.32 | 0.00239 |
| GO:0031981 | nuclear lumen | 40 | 8 | 2.58 | 0.00332 |
| GO:0043226 | organelle | 2048 | 151 | 131.92 | 0.00362 |
| GO:0043229 | intracellular organelle | 2035 | 150 | 131.08 | 0.00395 |

| | | | | | |
|---|---|---|---|---|---|
| GO:0000228 | nuclear chromosome | 6 | 3 | 0.39 | 0.00456 |
| GO:0005694 | chromosome | 9 | 3 | 0.58 | 0.01657 |
| GO:0031974 | membrane-enclosed lumen | 53 | 8 | 3.41 | 0.01862 |

Supplemental Table 2: GO term enrichment for maize Pfam domains with high PPA (PPA > 40). Bolded p-values pass a 5% Bonferroni multiple testing correction. Only the top 30 GO terms are shown.

| GO ID | Term | Annotated | Significant | Expected | p-value |
|---|---|---|---|---|---|
| GO:0032991 | protein-containing complex | 157 | 47 | 7.73 | **1.00E-27** |
| GO:0043228 | non-membrane-bounded organelle | 237 | 44 | 11.66 | **1.10E-16** |
| GO:0043232 | intracellular non-membrane-bounded organelle | 237 | 44 | 11.66 | **1.10E-16** |
| GO:0033176 | proton-transporting V-type ATPase complex | 11 | 11 | 0.54 | **2.60E-15** |
| GO:0016469 | proton-transporting two-sector ATPase complex | 12 | 11 | 0.59 | **3.00E-14** |
| GO:0005694 | chromosome | 184 | 36 | 9.06 | **3.20E-14** |
| GO:0005667 | transcription regulator complex | 21 | 12 | 1.03 | **2.30E-11** |
| GO:0000221 | vacuolar proton-transporting V-type ATPase | 8 | 8 | 0.39 | **2.70E-11** |
| GO:0016471 | vacuolar proton-transporting V-type ATPase | 8 | 8 | 0.39 | **2.70E-11** |
| GO:0033178 | proton-transporting two-sector ATPase complex | 8 | 8 | 0.39 | **2.70E-11** |
| GO:0033180 | proton-transporting V-type ATPase, V1 domain | 8 | 8 | 0.39 | **2.70E-11** |
| GO:0000126 | transcription factor TFIIIB complex | 12 | 9 | 0.59 | **2.40E-10** |
| GO:0090576 | RNA polymerase III transcription regulation | 13 | 9 | 0.64 | **7.60E-10** |

| | | | | | |
|---|---|---|---|---|---|
| GO:0098796 | membrane protein complex | 22 | 11 | 1.08 | **1.10E-09** |
| GO:0098687 | chromosomal region | 34 | 12 | 1.67 | **2.50E-08** |
| GO:0000228 | nuclear chromosome | 120 | 22 | 5.91 | **2.80E-08** |
| GO:1902494 | catalytic complex | 98 | 19 | 4.82 | 1.10E-07 |
| GO:0031981 | nuclear lumen | 226 | 30 | 11.12 | 1.50E-07 |
| GO:0031974 | membrane-enclosed lumen | 244 | 30 | 12.01 | 8.30E-07 |
| GO:0043233 | organelle lumen | 244 | 30 | 12.01 | 8.30E-07 |
| GO:0070013 | intracellular organelle lumen | 244 | 30 | 12.01 | 8.30E-07 |
| GO:0000781 | chromosome, telomeric region | 8 | 5 | 0.39 | 1.30E-05 |
| GO:0000428 | DNA-directed RNA polymerase complex | 7 | 4 | 0.34 | 0.00017 |
| GO:0030880 | RNA polymerase complex | 7 | 4 | 0.34 | 0.00017 |
| GO:0061695 | transferase complex, transferring phosphatase | 7 | 4 | 0.34 | 0.00017 |
| GO:0000775 | chromosome, centromeric region | 26 | 7 | 1.28 | 0.00018 |
| GO:0000178 | exosome (RNase complex) | 16 | 5 | 0.79 | 0.00075 |
| GO:1905354 | exoribonuclease complex | 16 | 5 | 0.79 | 0.00075 |
| GO:0005774 | vacuolar membrane | 45 | 8 | 2.21 | 0.00126 |
